# The organization of leukotriene biosynthesis on the nuclear envelope revealed by single molecule localization microscopy and computational analyses

**DOI:** 10.1101/521625

**Authors:** Angela B. Schmider, Melissa Vaught, Nicholas C. Bauer, Hunter L. Elliott, Matthew D. Godin, Giorgianna E. Ellis, Peter Nigrovic, Roy J. Soberman

## Abstract

The initial steps in the synthesis of leukotrienes are the translocation of 5-lipoxygenase (5-LO) to the nuclear envelope and its subsequent association with its scaffold protein 5-lipoxygenase-activating protein (FLAP). A major gap in our understanding of this process is the knowledge of how the organization of 5-LO and FLAP on the nuclear envelope regulates leukotriene synthesis. We combined single molecule localization microscopy with Clus-DoC cluster analysis, and also a novel unbiased cluster analysis to analyze changes in the relationships between 5-LO and FLAP in response to activation of RBL-2H3 cells to generate leukotriene C_4_. We identified the time-dependent reorganization of both 5-LO and FLAP into higher-order assemblies or clusters in response to cell activation via the IgE receptor. Clus-DoC analysis identified a subset of these clusters with a high degree of interaction between 5-LO and FLAP that specifically correlates with the time course of LTC_4_ synthesis, strongly suggesting their role in the initiation of leukotriene biosynthesis.

## Introduction

All cells must integrate and transduce multiple extracellular signals to achieve an appropriate functional response. In mast cells, the classic pathway of cell activation in response to an allergen is initiated by antigen binding to allergen-specific IgE antibodies coating mast cells via the IgE receptor (FcεR1; FcERI; UniProtKB: P12319) [1-3]. Antigen binding triggers the aggregation of FcεR1 receptors, activating their downstream pathways by recruiting a series of kinases to the cytoplasmic tail, and prolongs its presence on the plasma membrane [2, 4-7]. The kinase cascade sets in motion multiple processes including degranulation and the synthesis of leukotriene B_4_ (LTB_4_) [8, 9] and LTC_4_ [10, 11]. Activated mast cells predominantly make LTC_4_ [10, 11] and to a lesser extent LTB_4_ [9, 12]. In all cases, cells must generate a balanced response of lipid mediators and gene expression that is proportional to the strength, duration, and composition of the stimuli.

One way cells integrate signals is by the organization and disassembly of higher order multiprotein assemblies; especially those regulated by transient weak interactions [13, 14]. These structures must be assembled at the right time with the correct spatial localization. Modulating their composition and organization by orchestrating changes in the relationships between member proteins, such as by post-translational modification, can determine cellular responses to stimuli [13-15].

Leukotrienes (LTs), arachidonate 5-lipoxygenase (5-LO; UniProtKB: P09917) products of arachidonic acid (AA) metabolism, play a major role in initiating and amplifying inflammatory diseases, ranging from asthma to cardiovascular disease. Because of the dire consequences of the unregulated activation of this pathway, cells have evolved a series of complex control mechanisms to prevent the inadvertent initiation of LT synthesis. One strategy is based on the spatial segregation of the biosynthetic enzymes in different cellular compartments. In resting cells, 5-LO is localized in the nucleus and cytosol, and cytosolic phospholipase A_2_ (cPLA_2_; UniProtKB: P47712) is in the cytosol [16-21]. In activated mast cells, calcium influx triggers translocation of cPLA_2_ to the Golgi and ER membranes [16, 17], and of 5-LO to the nuclear envelope [19, 21], a main site LT synthesis. cPLA_2_ releases AA from the membrane phospholipids. To form the core of the LT synthetic complex on the nuclear envelope, [22, 23], AA associates with the homotrimeric integral membrane scaffold protein, arachidonate 5-lipoxygenase-activating protein (FLAP; UniProtKB: P20292). This event alters the relationship between the N-and C-terminal domains of FLAP [23] leading to recruitment of membrane-associated 5-LO [22, 23]. Though this is a transient, weak interaction requiring chemical crosslinking to identify biochemically [24], the 5-LO–FLAP complex functions to efficiently present AA to 5-LO and initiate synthesis of the parent LT, LTA_4_. In the presence of LTC_4_ synthase LTA_4_ is converted to LTC_4_, the first of the cysteinyl LTs[25] and is carried out of the cell by multidrug resistance-associated protein 4 (ABCC4; MRP4; UniProtKB: O15439) [26].

The formation and organization of higher order assemblies of receptors and enzymes is now considered an important regulatory process in signaling [27, 28]. This principle has been established for inflammasomes and signalosomes [29, 30]. In the context of the 5-LO pathway, the organization of the core LT synthetic complex into higher order assemblies could have several potential benefits. First, it could buffer the system from an “all or nothing” response to individual/random signals. Second, since AA diffuses through membranes and is rapidly re-esterified (28), it wcould provide a mechanism to increase the local concentration of the substrate and facilitate its availability to FLAP. Finally, it would provide a platform to integrate multiple signals.

We combined two-color direct stochastic optical reconstruction microscopy (dSTORM) single molecule localization microscopy (SMLM) with Clus-DoC analysis [31], and also linked single-color (conventional) STORM with a second unbiased clustering algorithm to define the time-dependent assembly and disassembly of higher order organizations of 5-LO and FLAP on the nuclear envelope. We identified a subset of clusters that contained a high degree of interaction between 5-LO and FLAP. Their assembly and disassembly was directly correlated with the synthesis of LTC_4_.

## Results

### Analysis workflow for developing a model of 5-LO and FLAP on the nuclear envelope

Analysis of the structure and characteristics of higher order assemblies has been facilitated by recent SMLM superresolution techniques combined with computational approaches. To determine 5-LO and FLAP organization on the nuclear envelope, two different computational analyses were applied following image acquisition (Fig 1). Tab-delimited text files (.txt) containing the localization list from each image set were output for each approach. We employed a clustering and colocalization algorithm, Clus-DoC [31] to two-color dSTORM data (purple). Clus-DoC assigns a degree of colocalization (DoC) score to each localization and determines percent colocalization, as well as cluster properties including the number of clusters in a region of interest (ROI), cluster area, and density. We also calculated other parameters such as the percent of interacting localizations inside clusters. In a second, supportive approach, we developed a novel unbiased clustering analysis based on a variable bandwidth mean-shift algorithm for conventional STORM data (orange). This analysis makes no assumptions about cluster size and is especially valuable for clusters curved around a membrane (S1 Fig).

**Fig 1.**
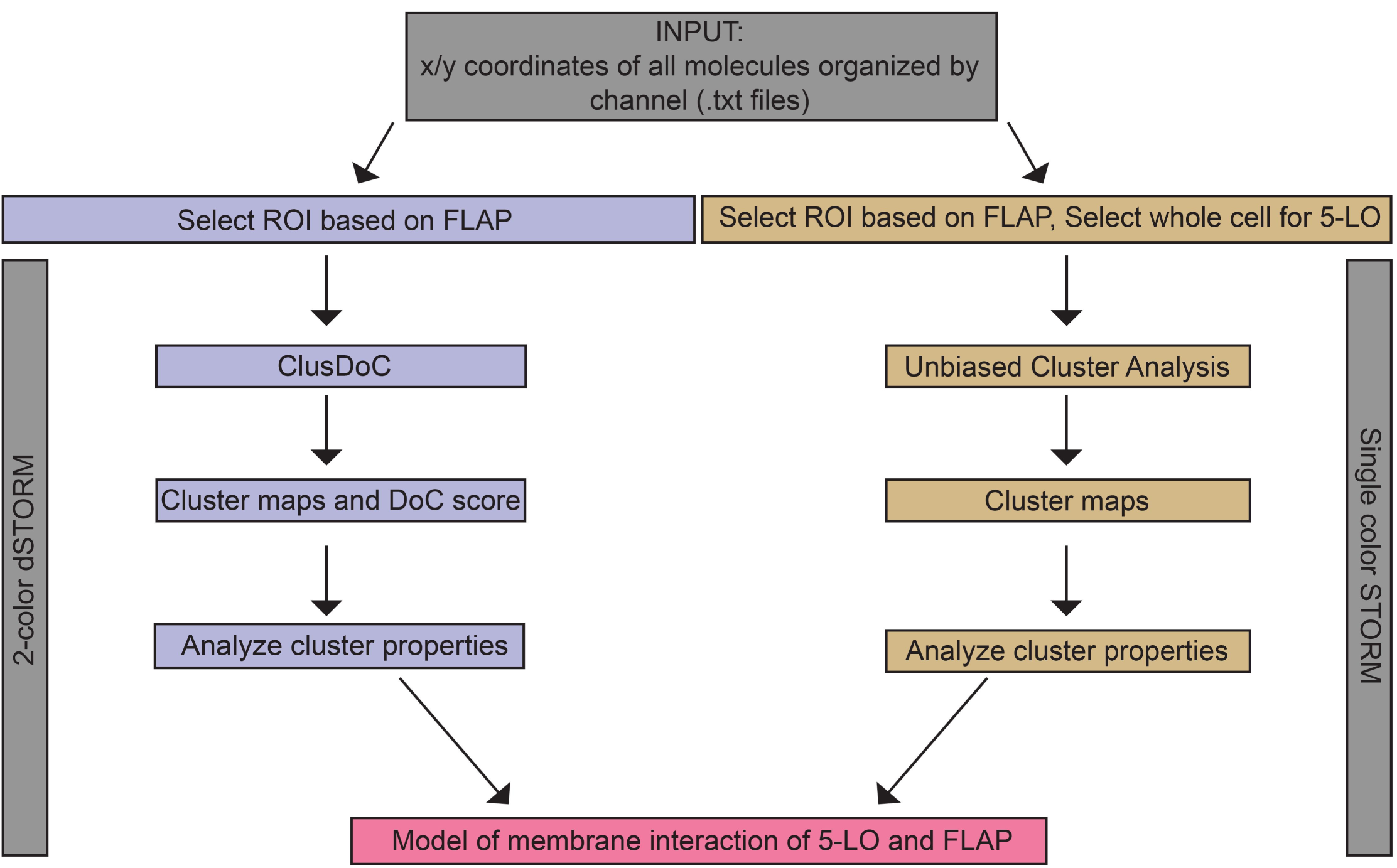
Analysis workflow for measuring the properties of 5-LO and FLAP and their relationship on the nuclear envelope. Initially, a .txt file with tab-delimited x/y coordinates of all localizations from both channels was generated. To analyze two-color dSTORM (purple), FLAP localizations were used to define a region of interest (ROI) around the nuclear envelope, which was also applied to 5-LO within the Clus-DoC user interface. Degree of colocalization (DoC) scores were calculated for each localization. DBSCAN detected clusters and defines cluster contours, yielding cluster maps and cluster properties. For conventional STORM (orange), FLAP localizations were used to define an ROI around the nuclear envelope, or in the perinuclear region and nucleus for 5-LO. Within unbiased cluster analysis, cluster maps with cluster properties are determined. Data from both methods were combined to produce a model of membrane reorganization of 5-LO and FLAP.

### Relationship of 5-LO and FLAP on the nuclear envelope

RBL-2H3 cells were primed with anti-TNP IgE and then stimulated with TNP-BSA for 2, 5, 7, and 10 min. Total media concentrations of LTC_4_ at each time point were measured, with significant LTC_4_ accumulating by 5 min and reaching peak levels by 7 min (Fig 2A). Cells were washed, fixed, and co-stained for 5-LO (ATTO 488; left/green) and FLAP (Alexa Fluor 647; middle/red), then imaged by dSTORM (Fig 2B). 5-LO is present at the nuclear membrane at 5 and 7 min (Fig 2B), whereas FLAP is present on the nuclear membrane at all time points (Fig 2B).

**Fig 2.**
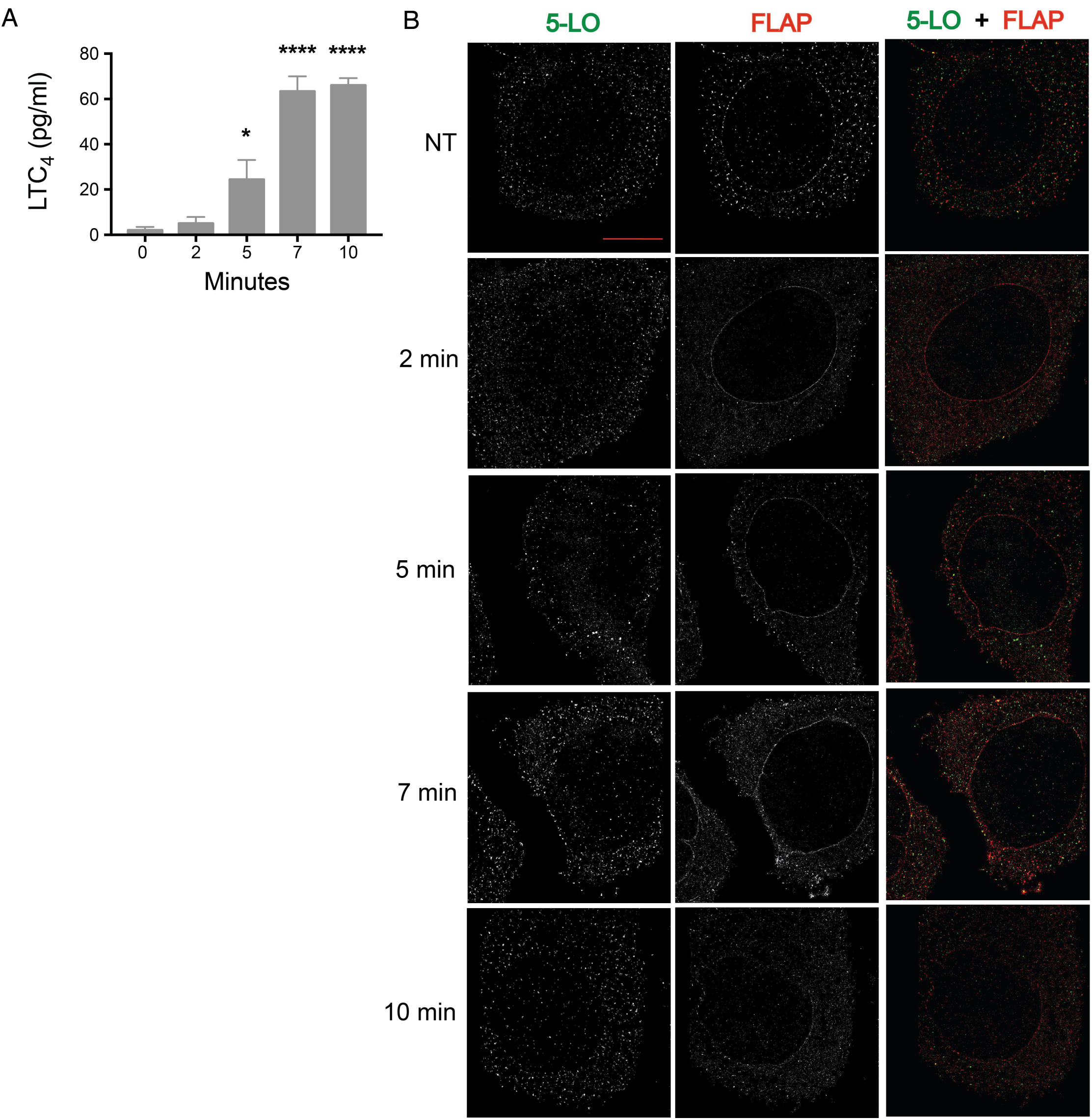
5-LO and FLAP localization on the nuclear envelope on mast cell activation by two-color dSTORM. RBL-2H3 cells (100,000 per well) were primed with anti-TNP IgE then activated with TNP-BSA for 0, 2, 5, 7, and 10 min, or left not treated (NT). Localization data was collected by two-color dSTORM. (A) Total media concentration of LTC_4_ (pg/mL) was measured by enzyme immunoassay. Mean ± SEM. Time points significantly different from 0 min (1-way ANOVA with Bonferroni post-hoc test) indicated by *p < 0.05, ****p < 0.0005. (B) Representative two-color STORM images of 5-LO (left panel, green) and FLAP (middle panel, red) following activation over time. Scale bar = 5 µm.

### 5-LO and FLAP colocalize on the nuclear envelope

Colocalization of 5-LO and FLAP in dSTORM images of activated mast cells at NT, and 2, 5, 7, and 10 min post-activation was determined using Clus-DoC [31]. Analysis was restricted to the nuclear envelope by manually drawn regions of interest (ROIs) based on FLAP localization. Fig 3A shows both the localization maps (left panels) and colocalization maps colored by DoC score, −1 (anticorrelated) to 1 (correlated) (right panels). The representative cells shown are identical to those shown in Fig 2B. The increase in orange and red localizations at 5 and 7 min represent increased colocalization between 5-LO and FLAP (Fig 3A, right panels). Fig 3B shows histograms of DoC scores in both directions (left: 5-LO to FLAP; right: FLAP to 5-LO) from the ROIs at NT (top) and 7 min (bottom). The frequency distributions for the ROIs selected at 2, 5 and 10 min are shown in S2 Fig. The percent of colocalized localizations, defined as the fraction of localizations with a DoC score ≥ 0.4, was determined for each ROI. At 5 and 7 min a two-fold increase in the percent of 5-LO molecules that were colocalized with FLAP was detected compared with NT (Fig 3D, left). No such difference was detected comparing FLAP to 5-LO due to high levels of FLAP compared with 5-LO (Fig 3D, right). These times correspond with maximal observed LTC_4_ synthesis (Fig 2A).

**Fig 3.**
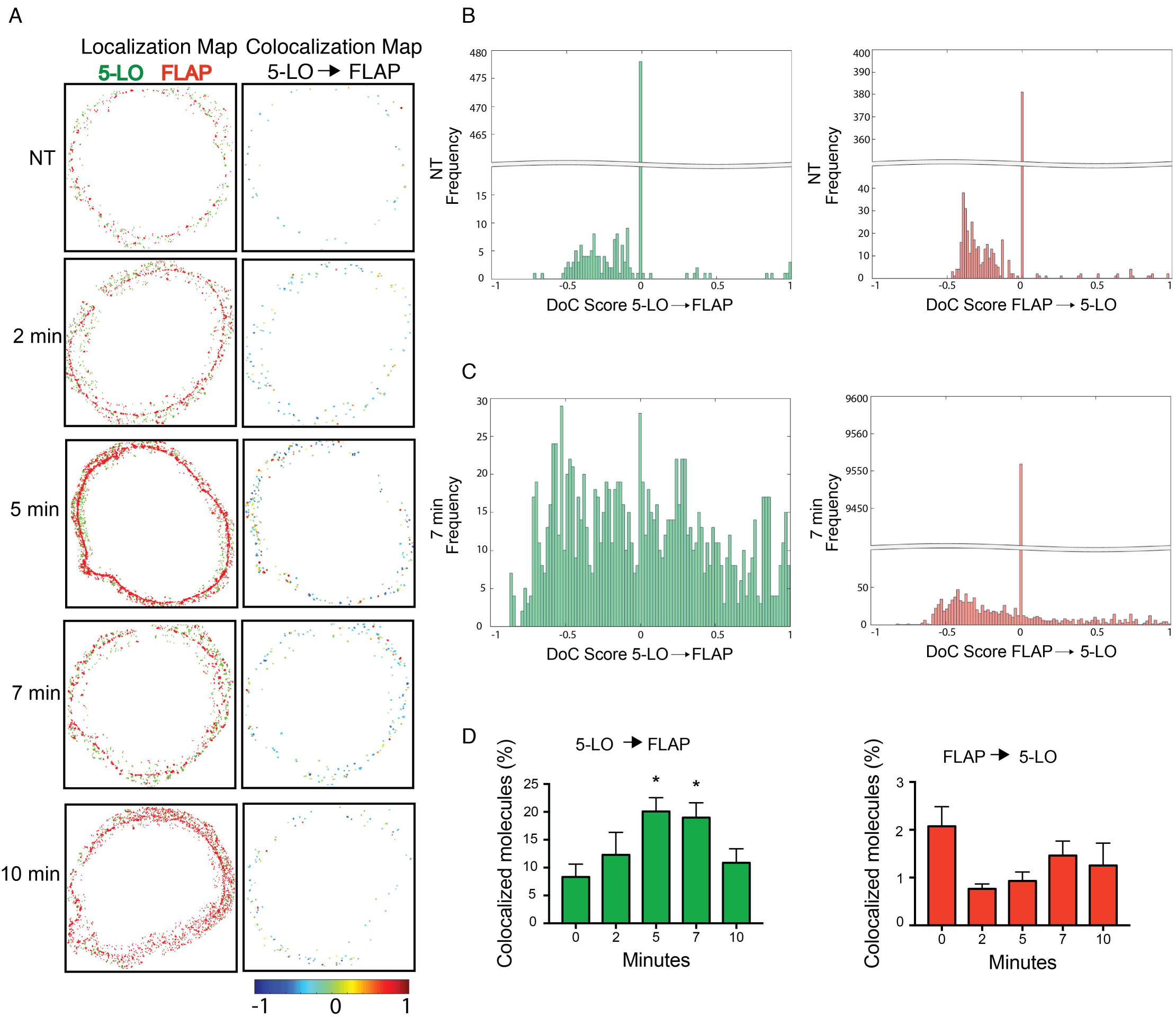
5-LO and FLAP colocalize on the nuclear envelope by two-color dSTORM. RBL-2H3 cells were primed with anti-TNP IgE then activated with TNP-BSA for 2, 5, 7 and 10 min, or left not treated (NT). Localization data was collected by two-color dSTORM and analyzed with ClusDoC. Data shown is from the representative cells shown in Fig 2. (A) Localization maps for 5-LO (green) and FLAP (red) (left panels) and colocalization maps for 5-LO relative to FLAP. 5-LO molecules are color-coded according to their degree of colocalization (DoC) scores (right panels, score bar at the bottom). (B,C) Histograms of DoC scores of all molecules for 5-LO (green) and FLAP (red) from representative cells at (B) 0 and (C) 7 min. (D) Percent colocalization of 5-LO molecules with FLAP from all ROIs (left panel; green) and percent colocalization of FLAP molecules with 5-LO from all ROIs (right panel; red). Statistical significance was assessed by Kolmogorov-Smirnov test, with significance indicated by *p < 0.05. Bars show mean ± SEM from 4 to 21 cells over 3 separate experiments.

### The formation of higher order assemblies of 5LO and FLAP

If higher order assemblies of 5-LO and FLAP play a regulatory role in LT synthesis, their formation and disassembly on the nuclear envelope should be correlated with LTC_4_ production. We employed the Clus-DoC algorithm to test this hypothesis. The algorithm measured the colocalization of the localizations and then sorted them into clusters or excluded them as outliers. The algorithm then calculated the characteristics of the identified clusters, allowing changes in molecular organization to be detected. Cluster maps of 5-LO and FLAP for the cells analyzed in Figs 2-3 are shown in S3 Fig; cluster contours are indicated by the black outline and points outside of clusters are shown in gray. Two levels of cluster limits were applied as modifications of Clus-DoC. First, a threshold of >5 localizations for 5-LO or >10 localizations for FLAP were used to define a true cluster. A higher threshold was chosen for FLAP because it functions as a homotrimer [24], increasing the chance that identified clusters contain at least 3 functional FLAP molecules. A second level was used to distinguish clusters with no colocalization (no interaction clusters (NIC); white bars), a high degree of colocalization (high interaction clusters (HIC); black bars) denoting at least 5 localizations with a DoC score of ≥ 0.4, and all other clusters (low interaction clusters (LIC); gray bars). The properties of clusters were compared between HIC, LIC, and NIC at different time points (Fig 4). The number of HIC per ROI increased from 0.3 at NT and 2.3 at 2 min to 6.4 and 7.8 at 5 and 7 min, respectively, and returned to 0.6 at 10 min (Fig 4A). Because there were <3 ROIs containing HIC at NT, 2 and 10 min, we excluded them from further analysis. The appearance and disappearance of HIC correlates with peak LTC_4_ synthesis, suggesting that the formation of HIC is required for LTA_4_ synthesis and that disassembly of these clusters is a critical step in the termination of synthesis.

**Fig 4.**
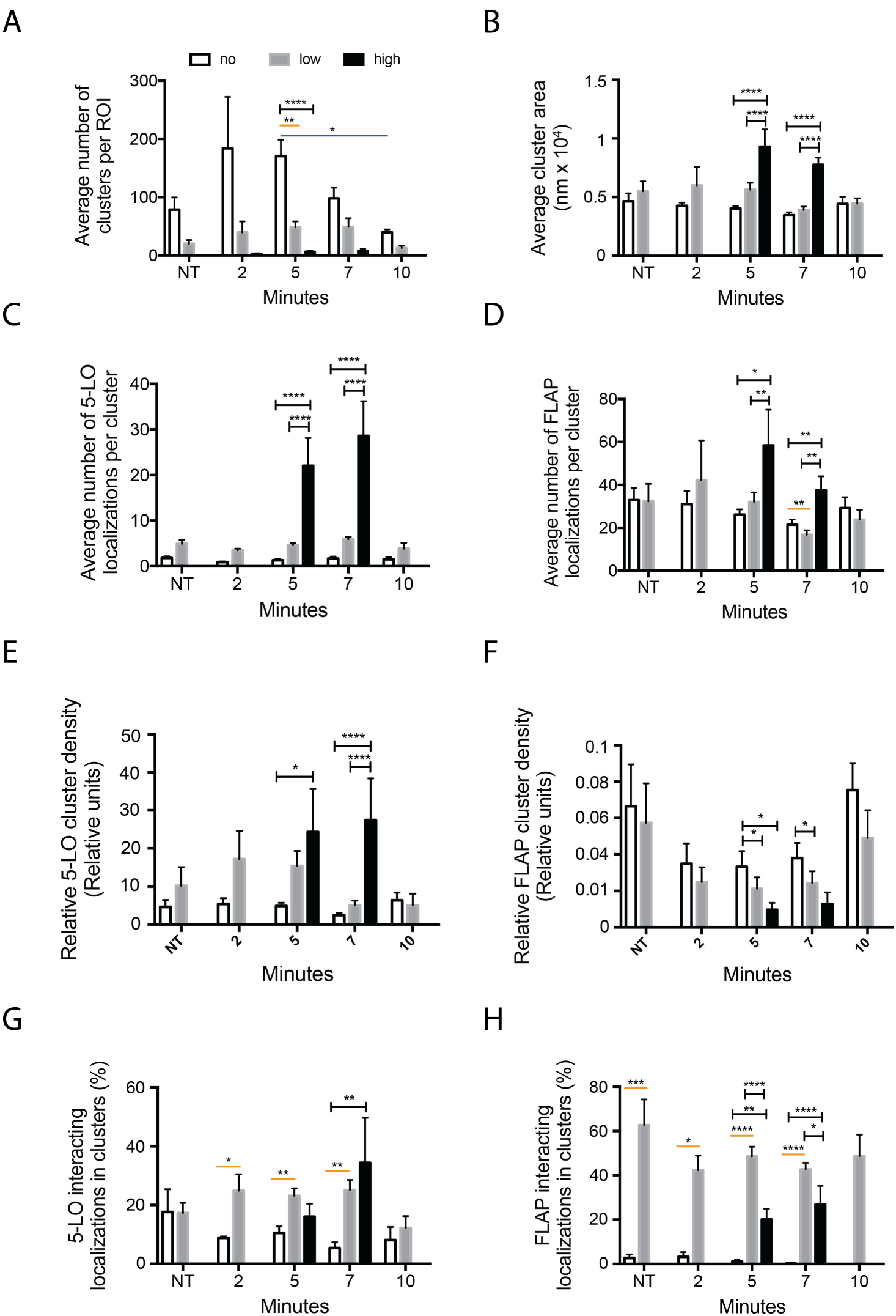
Activation changes 5-LO and FLAP cluster properties by two-color dSTORM. RBL-2H3 cells were primed with anti-TNP IgE then activated with TNP-BSA for 2, 5, 7, and 10 min, or left not treated (NT). Localization data was collected by two-color dSTORM and analyzed with Clus-DoC. Clusters were defined as having either ≥ 5 5-LO localizations or 10 FLAP localizations, and were split into 3 categories based on the number of founding localizations with a degree of colocalization (DoC) score ≥ 0.4 indicating interaction: no interaction clusters (NIC; white bars) with 0 interacting localizations; low interaction clusters (LIC; gray bars) with between 1 and 4 interacting localizations; and high interaction clusters (HIC; black bars) with 5 or more interacting localizations. (A) Average number of clusters per ROI. (B) Average cluster area. (C,D) Average number of 5-LO and FLAP localizations per cluster, respectively. (E,F) Relative density of 5-LO and FLAP in clusters, respectively. Relative density of clusters was calculated by dividing the local density within 20 nm of each localization by the average density of the cluster, a measure of the local concentration maxima within the cluster. (G,H) Percent of interacting 5-LO and FLAP localizations located in clusters, respectively. Time points for a cluster type with fewer than 3 values for that cluster type were excluded. One-way ANOVA with Tukey’s post-hoc multiple comparisons test was performed to determine significance among interaction groups where *p < 0.05, **p < 0.005 and ****p < 0.0001; within timepoint, HIC–NIC or HIC–LIC indicated by black brackets, LIC–NIC by orange line; between timepoints by blue line. Bars show mean ± SEM of clusters within 4 to 21 cells over 3 independent experiments.

### Properties of HIC link them to LT synthesis

Changes in six separate properties of HIC are temporally correlated with LTC_4_ synthesis. First, their area is approximately 2-fold larger than LIC and NIC at 5 and 10 min after activation, respectively (Fig 4B). Similarly, the average number of 5-LO localizations in HIC are 10-20 fold higher than those in NIC and LIC (Fig 4B). In contrast, number of 5-LO localizations in NIC and LIC remained unchanged over time (Fig 4C). The number of FLAP molecules in HIC was approximately 3-fold higher (5 min) and 2-fold higher (7 min) than in LIC and NIC, which did not change over the time course of the experiment (Fig 4D).

The relative density of 5-LO in clusters remained unchanged within NIC and LIC over time. Relative density is calculated as the average of local density within 20 nm of each localization in the cluster, divided by the average cluster density, providing a measure of the distribution of localizations within a cluster. However, the relative density of 5-LO in HIC was elevated 2-fold at 5 min and 15-fold greater than that of NIC and LIC at 7 min (Fig 4E). Interestingly, FLAP relative densities for NIC were 2-fold lower at 5 and 7 min than in LIC or HIC (Fig 4F).

We next calculated the percent of all 5-LO and FLAP interacting localizations that are in clusters. At 2, 5 and 7 min after activation, the vast majority of 5-LO molecules were found in either LIC or HIC (Fig 4F). This was true for FLAP at all time points (Fig 4G). Taken together, these data link the formation and disassembly of HIC to LTC_4_ formation.

### The dynamic organization of 5-LO analyzed by conventional STORM

Conventional STORM experiments in combination with unbiased cluster analysis revealed similar cluster properties to those revealed using Clus-DoC. The cells were prepared identically but stained only for 5-LO and imaged by STORM. Because conventional STORM was used in these experiments, FLAP localizations were not available to define the nucleus in the same cell. Therefore, the whole nucleus and perinuclear area was included in the ROI. RBL-2H3 cells were analyzed at 0, 2, 5, and 10 min after cell priming and activation. 5-LO localizations in representative cells acquired by STORM are shown in Fig 5A (top panels). Unbiased cluster analysis grouped localizations into clusters of >3 localizations. The organization of 5-LO was not uniform across the nucleus and the region that included the nuclear envelope, both in size and shape (Fig 5A, middle panels) and number of localizations per cluster (Fig 5A, lower panels, pseudocolor legend below). Convex hulls represent cluster contour (Fig 5A, middle panels). An outline derived from immunofluorescence illustrates where the nuclear envelope lies. Unbiased cluster analysis showed that both the number of 5-LO clusters per cell and the number of localizations per cluster increased at 2 and 5 min post-activation compared to control and decreased to baseline at 10 min post-activation (Fig 5A, lower panels).

**Fig 5.**
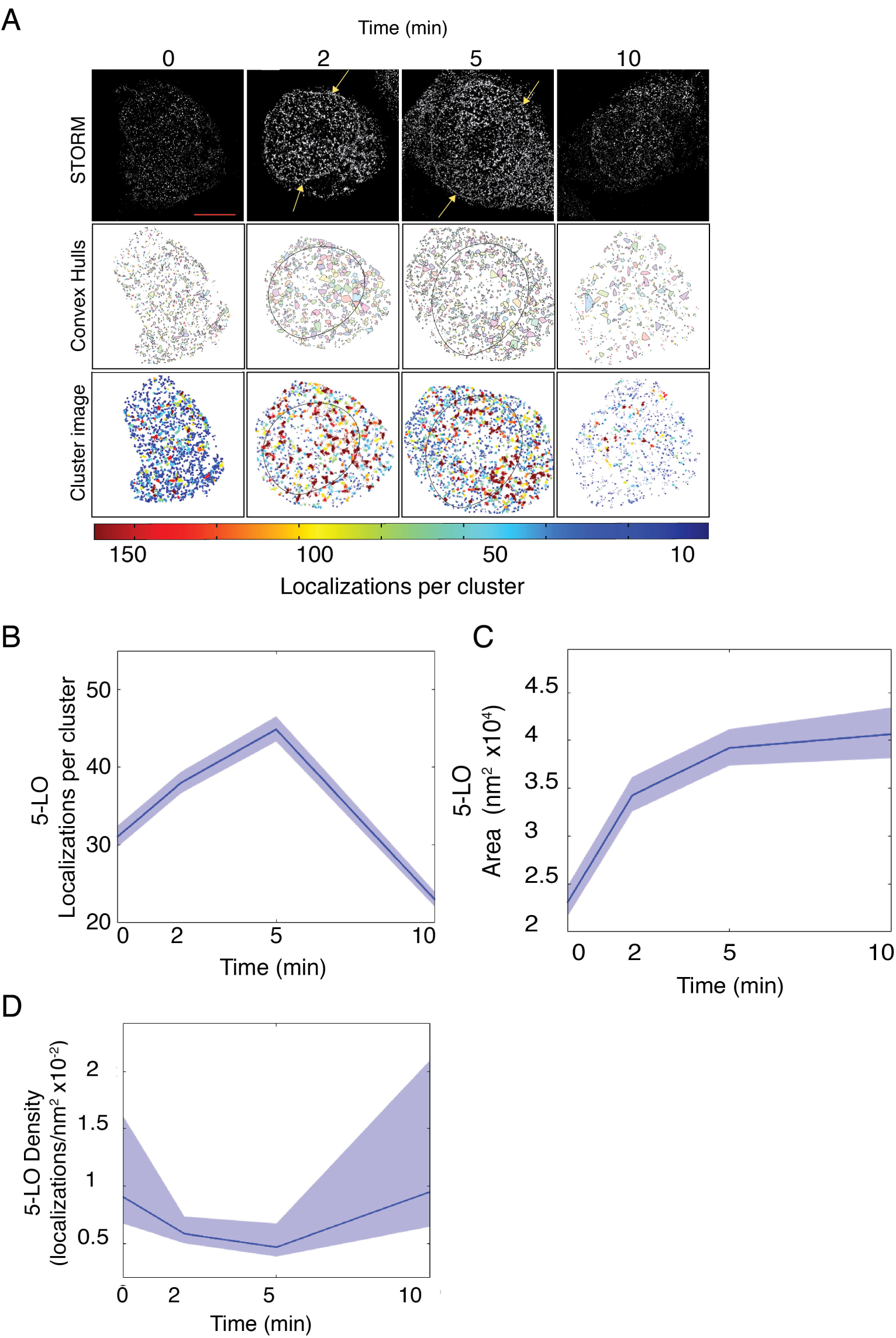
Activation organizes 5-LO into clusters by conventional STORM. RBL-2H3 cells were primed with anti-TNP IgE then activated with TNP-BSA for 0, 2, 5, and 10 min and analyzed as shown S1 Fig. Cells were imaged with conventional STORM and cluster properties were analyzed with unbiased cluster analysis. (A) Detailed STORM analysis of 5-LO following priming and activation over time. Grayscale STORM images show localizations (upper panels), clusters are shown as convex hulls (middle panels, arbitrary colors) and 5-LO clusters are shown as points colored by number of localizations in the cluster (lower panels, legend bar below). (B) The average number of localizations per cluster over time. (C) The average area of 5-LO clusters over time. (D) The average density of 5-LO clusters over time. Plots show mean ± 95% confidence intervals of all pooled clusters identified in cells for that time point. At least 3 separate experiments collected between 10 and 30 cells. Scale bar = 5 μm.

Using 95% confidence intervals (CI), the mean number of localizations associated with clusters (pooled across all cells of a condition and time point) of 5-LO increased from 32 to 38, 2 min after the addition of antigen and peaked at 5 min, having 45 localizations per cluster (a 25% increase). After 10 min, the mean number of localizations dwindled to 22, which is lower than unstimulated conditions (0 min), suggesting complex disassembly (Fig 5B). The mean size of 5-LO cluster areas was also analyzed using 95% CIs. In non-activated cells, there were small numbers of 5-LO associated with the nuclear envelope and a correspondingly low number of localizations per cluster were detected in unstimulated conditions. The mean area of 5-LO clusters identified on the nuclear envelope was ~2.1×10^4^ nm^2^ (Fig 5C). At 2 min following cell activation, the mean cluster area climbed to 3.4×10^4^ nm^2^ and then to 4×10^4^ nm^2^ at 5 min and 10 min (Fig 5C). The mean cluster density declined over time, decreasing by 50% at 5 min and returning to control levels at 10 min post-activation (Fig 5D).

To determine whether 5-LO clusters of a specific size, area, or density are assembled after cell activation, we employed weight-normalized histograms of the ROIs in Fig 5. There was a progressive increase in the frequency of the clusters containing the highest numbers of localizations (greater than 250 localizations or more) at 2 and 5 min stimulation, then decreasing at 10 min (S4 Fig). The organizational shift from a high frequency of clusters in control cells containing few localizations per cluster (less than 100) to significantly more localizations per cluster (greater than 250) occurred at 2 and 5 min after cell activation, concurrent with LT production (Fig 2A). Stimulating cells for 10 min resulted in the same pattern of localization frequency as control. The mean number of 5-LO localizations per cluster was discernably higher following cell activation at 2 and 5 min compared to control (S4 Fig, inset). Following stimulation, the frequency of clusters with larger areas increased over time (S4 Fig, inset), while the densities of clusters remained constant (S4 Fig, inset).

The formation of higher order FLAP assemblies was detected using conventional STORM in combination with unbiased cluster analysis. FLAP is primarily localized to the nuclear envelope and to a lesser extent on the ER membrane. Because we were interested in analyzing FLAP distribution in its primary localization, we used FLAP localizations to define the nuclear envelope for unbiased cluster analysis. We first probed whether FLAP would reorganize within the nuclear envelope following IgE priming and antigen activation. RBL-2H3 cells were primed and activated for 0 or 7 min. Fig 6A shows the STORM and cluster analyses images with convex hulls (middle panels) and numbers per cluster (lower panels, pseudocolor legend below). Weight-normalized histograms show that distribution of clusters shifts to the left after addition of antigen, whereas cells only primed with IgE contain a higher frequency of clusters with more localizations (Fig 6B). There was an increase in clusters containing between 250 and 450 localizations corresponding to ~83-150 trimers when cells were primed and activated (Fig 6B). A similar pattern was observed for cluster areas. Cells that were only primed with IgE contained a greater number of clusters with larger cluster areas compared to cells that were primed and activated with antigen (Fig 6C). The frequency profile for cluster density was marginally different between IgE-primed and IgE-antigen activated cells (Fig 6D). The mean number of localizations per cluster decreased approximately 25% following addition of antigen (Fig 6B, inset) and the mean cluster area decreased from 1.4×10^5^ nm^2^ to 0.8×10^5^ nm^2^ following antigen stimulation (Fig 6C, inset). In parallel, the mean density of clusters increased ~25% from 1.7×10^-3^ localizations/nm^2^ to ~2.4×10^-3^ localizations/nm^2^ after antigen stimulation, suggesting that following antigen, FLAP clusters become more compact.

**Fig 6.**
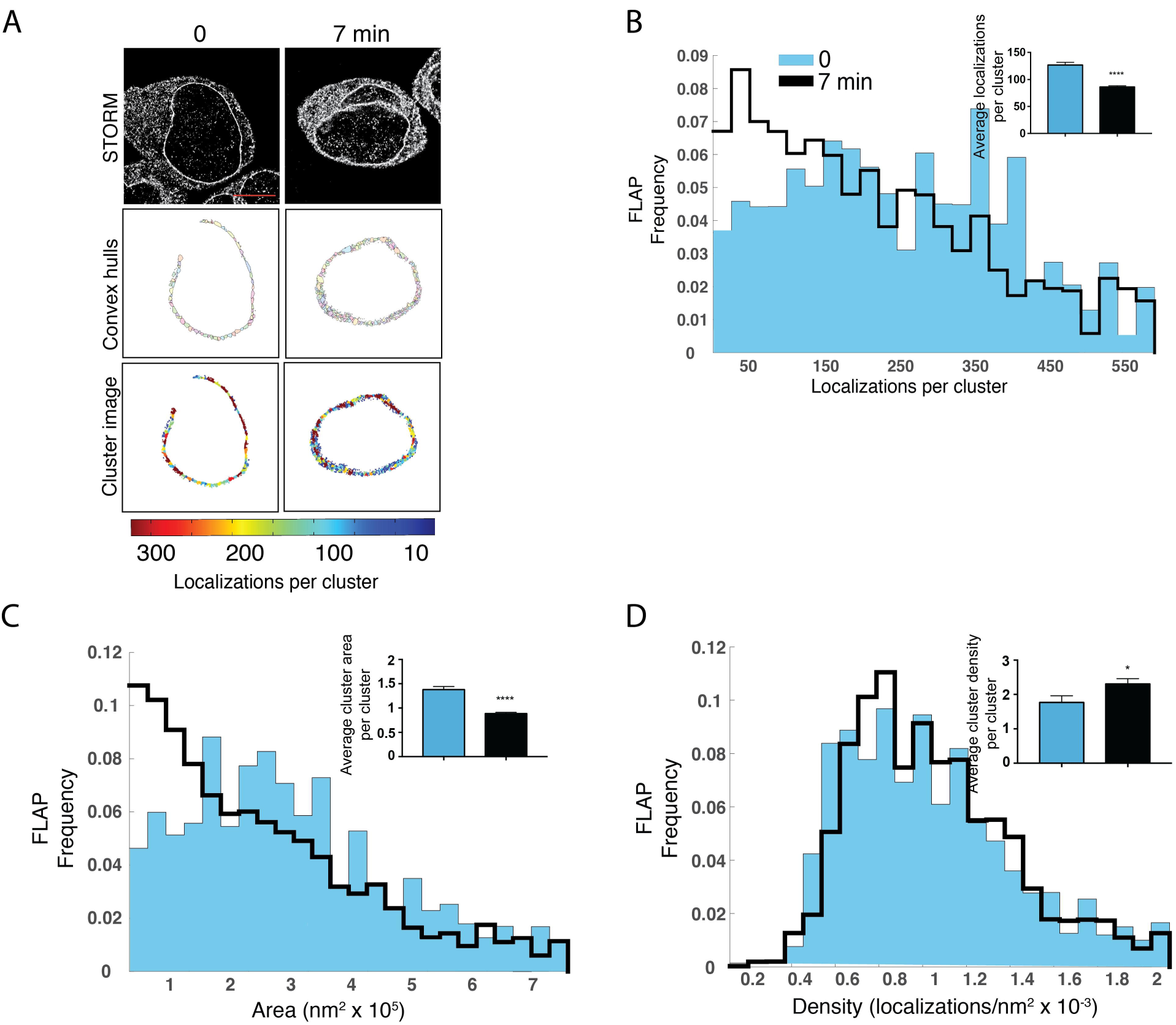
FLAP re-organization on the nuclear envelope. RBL-2H3 cells were primed with anti-TNP IgE then activated with TNP-BSA for 0 and 7 min then analyzed as shown Fig 1. Cells were imaged with conventional STORM and cluster properties were analyzed with unbiased cluster analysis. (A) Detailed STORM analysis of FLAP following priming and activation over time. Grayscale images show localizations (upper panels), cluster areas are shown as convex hulls (middle panels, arbitrary colors) and FLAP clusters are shown as points colored by number of localizations in the cluster (lower panels, legend bar below). Scale bar = 5 μm. (B-D) Normalized point-weighted histograms with inset bars showing mean ± SEM for (B) number of localizations, (C) cluster areas and (D) cluster densities. Student’s t-test was used to determine significance indicated by *p < 0.05 and ****p < 0.0005. At least 3 separate experiments collected between 10 and 30 cells.

Since the interaction of 5-LO with FLAP is dependent on the association of AA with FLAP [24], we tested whether the organization of 5-LO into higher order assembles was also dependent on AA (S5 Fig). Cells were primed with IgE and then stimulated by the addition of antigen for 7 min in the presence or absence of the cPLA_2_ inhibitor (Inh), or the presence or absence of the FLAP inhibitor MK886. When the release of AA was blocked (S5 Fig A-C, insets), there was no change in distributions for the number, area or density cluster properties of 5-LO localizations (S5 Fig, A-C, insets). As shown in weight-normalized histograms, the frequency of these clusters was unchanged (S5 Fig, A-C). When MK886 was added the properties of clusters changed. The mean number of localizations per cluster and cluster area both increased (S5 Fig, A, B insets), while the cluster density decreased (S5 Fig, inset). There was a higher frequency of clusters containing 100 or more localizations per cluster in cells pre-treated with MK886 compared to primed and activated cells without inhibitor (S5 Fig). A similar pattern was seen in cluster area where the frequency of clusters with larger areas was higher in cells pre-treated with MK886 (S5 Fig). Though the mean density per cluster decreased, the distribution was similar to that with cells treated with IgE and antigen alone and cells pre-treated with MK886 (S5 Fig).

In the presence of the cPLA_2_ Inh or the FLAP Inh following priming with IgE and antigen, the number of FLAP localizations per cluster decreased (Figures S5, inset). The shifts in the frequency distribution of clusters for each inhibitor were similar, with fewer localizations in each cluster compared to cells only primed and activated (S5 Fig). The mean cluster area slightly decreased when cells were exposed to the cPLA_2_ inhibitor during priming and antigen activation (S5 Fig, inset). MK886 had no effect on FLAP cluster area following cell priming and antigen activation (S5 Fig). Localization frequencies were left shifted when cells were exposed to either cPLA_2_ inhibitor or FLAP inhibitor. This was a slight, but notable decrease in the frequency of clusters containing 300 localizations or more (Fig 4D). Cluster density was unchanged in the presence of cPLA_2_ or FLAP inhibitors (S5 Fig).

## Discussion

Analysis of two color dSTORM data using the Clus-DoC algorithm allowed us to define three classes of clusters: NIC, LIC, and HIC, and provided strong evidence linking HIC to the synthesis of LTs (Figs 1-4). The peak changes in the number of HIC and their properties occurred at 5 and 7 minutes, times of maximal LTC_4_ generation. These changes include increases in area, the average number of 5-LO and FLAP localizations per cluster, and the percent of 5-LO and FLAP in HIC interacting with each other (Fig 4). As with 5-LO, the number of FLAP localizations in HIC increased at 5 and 7 min. Furthermore, almost all FLAP molecules that interact are in LIC or HIC. Interestingly, compared to LIC, the relative density of FLAP in HIC is lower. Using conventional STORM, the same trends were observed for the area and number of 5-LO localizations, whereas there was a modest decrease in average density (localizations per unit area) at 2 and 5 min post-activation, which may be attributed to the use of the whole nucleus as an ROI. The FLAP localization data obtained with conventional STORM identifies global changes in properties of clusters, but highlights the benefits of two-color dSTORM paired with clustering algorithms that consider both molecules of interest in characterizing higher order assemblies. Despite the limitations of conventional STORM, a role of FLAP in regulating higher order assemblies of 5-LO is supported by experiments with MK886, which occupies the extended AA binding site, in which the number of 5-LO molecules and area of 5-LO in clusters is increased. In contrast, cPLA_2_ Inh had no effect on organization, suggesting that overall re-organization of 5-LO or FLAP is independent of AA.

The number of localizations can be considered an estimate of the number of molecules in a cluster. Determining the absolute number of molecules from STORM data is difficult due to the stochastic blinking behavior of the fluorophores [32], but relative assessments should be less affected. In two-color dSTORM experiments the number of localizations per HIC was 20-30 for 5-LO and 40-60 for FLAP at 5 and 7 min. Because FLAP functions as a trimer [24], this data provides a rough estimate of a near-stoichiometric 5-LO:FLAP ratio of 1:1 to 2:1 in HIC. At 5 and 7 min post-activation, the peak of LT synthesis, a significant fraction of interacting localizations for both 5-LO and FLAP are localized in HIC. Together, these data strongly indicate a role for clustering in mediating LT synthesis.

The critical regulatory role for higher order assemblies, typified by signalosomes, is the new paradigm which must be considered when evaluating all membrane signaling processes [26-30]. These studies were based upon data obtained with crystallography and electron microscopy (EM) [21]. When combined with algorithms such as Clus-DoC, two-color dSTORM is a nanoscale SMLM approach that provides the ability to test for higher order organization while maintaining cellular organization. SMLM combined with cluster analysis has allowed the identification of the segregation of clusters of active and inactive integrin molecules on membranes [33]. iPALM has successfully elucidated the role of integrin configuration during adhesion [34]. Previous work by our laboratory using fluorescence lifetime imaging microscopy and biochemical crosslinking identified the interaction of 5-LO and FLAP [22, 23, 35]. This work was supported by subsequent studies, including those using overexpression approaches and/or non-physiological stimuli to probe the relationship between the two molecules [36-38]. None of these earlier approaches have addressed the critical concept of membrane organization at nanoscale and were limited by the technologies employed.

Overall, our work links higher order assemblies of 5-LO and FLAP to the initiation of LT synthesis on the nuclear envelope. Fig 7 shows our proposed model: At steady state (NT), small numbers of clusters (NIC; light gray outline) of 5-LO (green) or FLAP (red) exist on the nuclear membrane, few with 5-LO associated with FLAP (LIC; dark gray outline). By 2 min after activation, there is an increase in both total number of 5-LO and FLAP in NIC and LIC. Between 5 and 7 min post-activation, clusters with extensive interaction between 5-LO and FLAP (HIC; black outline) have formed. By 10 min, no HIC remain. Our work illustrates the power of using two-color dSTORM combined with algorithms such as Clus-DoC to analyze data, allowing us to unveil relationships not obtainable by conventional STORM and cluster analysis. Characterizing the mechanisms of higher order assembly and disassembly of biosynthetic complexes is ultimately essential to understanding the initiation of LT synthesis and the generation of other eicosanoids and products of eicosapentaenoic acid and docosahexanoic acid.

**Fig 7.**
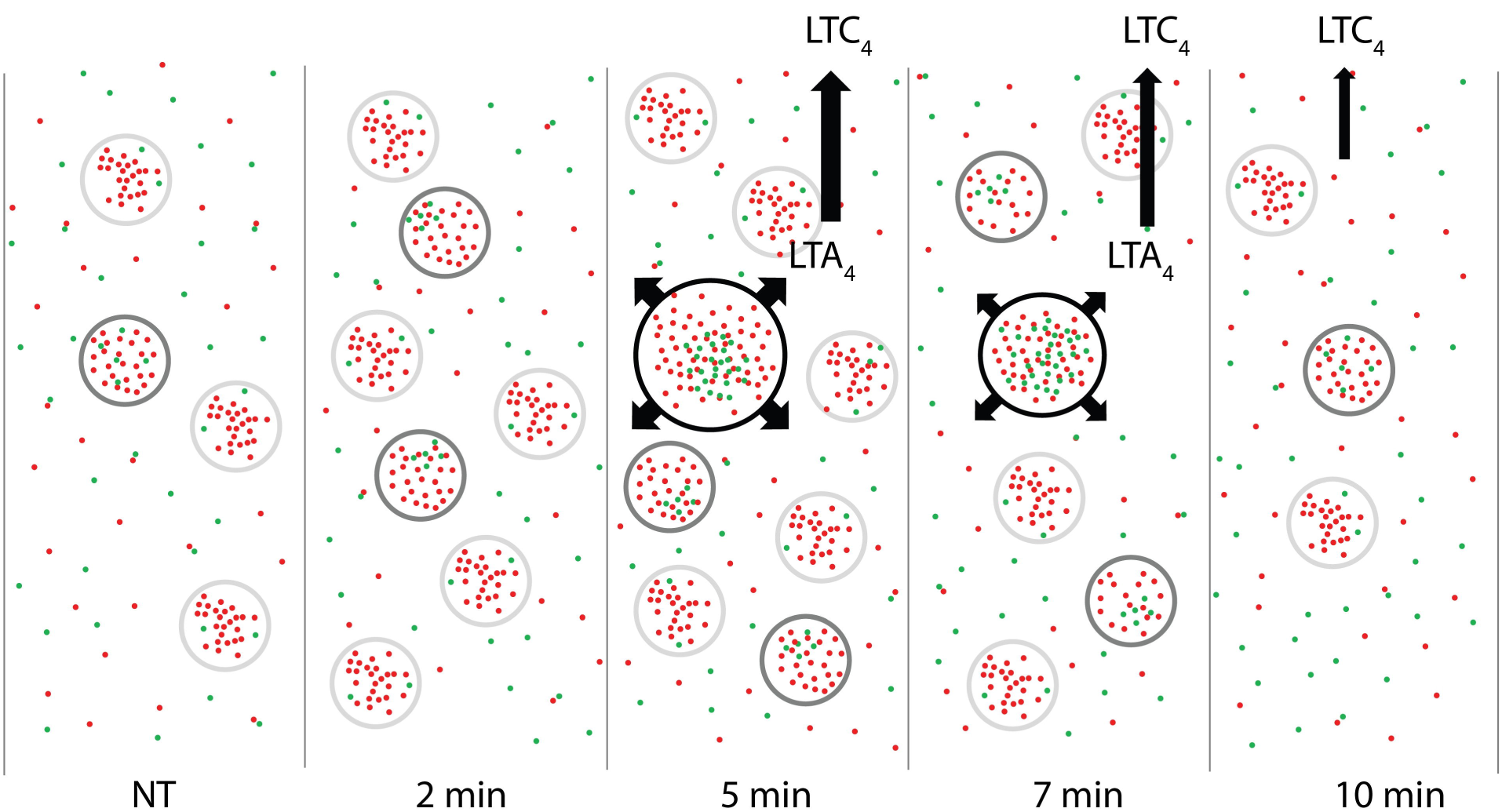
Proposed model linking HIC to LTC_4_ synthesis. At steady state (NT), small clusters (NIC, light gray outline) of 5-LO (green) or FLAP (red) exist on the nuclear membrane, few with 5-LO associated with FLAP (LIC, dark gray outline). By 2 min after activation, 5-LO and FLAP increasingly interact and more LIC are observed. Between 5 and 7 min after activation, large clusters with extensive interaction between 5-LO and FLAP have formed (HIC, black outline) and produce LTA_4_, which is converted to LTC_4_ then released from the cell. HIC are no longer observed at 10 min.

## Materials and methods

### Cell culture, activation, fixation, staining

RBL-2H3 cells (ATCC, CRL-2256) were maintained in Dulbecco’s modified Eagle’s medium (DMEM) supplemented with 10% heat-inactivated FCS, 2 mM glutamine, penicillin (100 units/mL), and streptomycin (0.1 mg/mL). The cells were primed with trinitrophenol (TNP)-specific IgE (0.1 μg/mL IgE-3 clone, BD Biosciences) at 37 °C for 45 min in Dulbecco’s phosphate buffered saline (DPBS) containing glucose, Ca^2+^, Mg^2+^ (Cellgro), and 0.1% BSA (DPBS/BSA) to load IgE onto FcεRI without LT synthesis. Following incubation with IgE, cells were washed with DPBS to remove unbound IgE. LTC_4_ synthesis was initiated by addition of TNP-conjugated BSA (TNP-BSA, 25 ng/mL in DPBS/BSA, Santa Cruz Biotechnology). (N-{(2S,4R)-4-(Biphenyl-2-ylmethyl-isobutyl-amino)-1-[2-(2,4-difluorobenzoyl)-benzoyl]-pyrrolidin-2-ylmethyl}8-3-(4-(2,4-dioxothiazoldin-5-ylidenemethyl)-phenyl]acrylamide (cPLA_2_ inhibitor, cPLA_2_ Inh, Calbiochem, EMD Biosciences), was used concomitantly with priming at 5 μM and MK-886 (FLAP Inh, Cayman Chemical) was also used concomitantly at 5 nM.

### LTC_4_ synthesis assay

To analyze LTC_4_ generation, an enzyme immunoassay kit for LTC_4_ (Cayman Chemical) was used per manufacturer’s instructions. Culture media was removed at desired times and stored at −80 °C until use.

### Stochastic optical reconstruction microscopy

#### Preparation of secondary antibodies

To produce activator-reporter tandem dye pair antibodies for conventional STORM, Donkey anti-rabbit and donkey anti-goat affinity purified secondary antibodies (H+L chains) (Jackson Laboratory) were conjugated to Cy3 (Jackson ImmunoResearch) (activator) and Alexa Fluor 647 (AF647, Life Technologies) (reporter). For the conjugation reactions, 65 μg of the secondary antibody was reacted with 3 μg of the activator dye (Cy3) and 1.5 μg of the reporter dye (AF647) in 103 mM carbonic buffer for 2 hours at room temperature in the dark.

To produce reporter antibodies for direct STORM, donkey anti-goat and donkey anti-mouse affinity purified secondary antibodies (H+L chains) (Jackson Laboratory) were conjugated to ATTO 488 (Millapore Sigma) or AF647 dyes, both with carboxylic acid succinimidyl ester moieties. For the conjugation reactions, 240 μg of the secondary antibody was reacted with 6 μg of dye in 56 mM carbonic buffer for 2 hours at room temperature.

After the reactions, the antibodies were separated from unconjugated dye by gravity filtration through Sephadex G-25 DNA grade size exclusion columns (GE Healthcare) by visual detection. Antibody and dye concentrations were determined using a NanoDrop spectrometer (Thermo Fisher) to record absorbance at 280 nm and at the dye absorbance peak. Antibody concentration was calculated by subtracting the contribution of each dye to absorbance at 280 nm using correction factors provided by the dye manufacturers (ATTO488: 0.1, Cy3B: 0.09, AF647: 0.03) and the molar extinction coefficient of the antibody (210,000).

The activator dye:antibody:reporter dye ratio for Cy3:donkey-anti-rabbit:AF647 was 0.8:1:2.1 with antibody concentration of 145 μg/μL. The ratio for conjugated Cy3:donkey-anti-goat:AF647 was 1.1:1:3.2 with antibody concentration of 140 μg/μL. The antibody:dye ratio for donkey-anti-mouse:ATTO488 was 1:2.7 with antibody concentration of 280 μg/μL. The ratio for donkey-anti-goat:AF647 was 1:1.7 with antibody concentration of 200 μg/μL.

#### Conventional STORM

After activation, cells were fixed and prepared for STORM as previously described [39]. The following day they were stained with antibodies to FLAP (Novus IMG 3160, 1:100) or to 5-LO (Santa Cruz H-120, sc-20785, RRID: AB_2226938, 1:20). Activator-reporter antibody was applied at 3 µg/mL for 1 h. Imaging buffer containing 147 mM βME and 1% (v/v) glucose oxidase with catalase (GLOX) was used to promote photoswitching and reduce photobleaching [39]. Cells were imaged in continuous mode on an inverted Nikon Ti-Eclipse STORM 3.0 system equipped with 100X/1.4 NA objective lens, iXon X3 EM CCD camera (Andor), and 647 nm (300 mW), 561 nm (150 mW) and 405 nm lasers (100 mW). 9,000 frames for each dye were collected at 30 ms exposure time. Localizations were identified with NIS Elements 3.0 (Nikon Instruments) and exported as tab-delimited text files.

#### Two-color dSTORM

After activation, cells were fixed and prepared for superresolution microscopy [39]. The following day they were probed with antibodies to FLAP (Novus, IMG 3160, 1:100) and to 5-LO (BD Biosciences, 610695, RRID: AB_398018). Secondary antibodies were applied at 3 µg/mL for 1 h. The imaging buffer containing 100 mM 2-mercaptoethanolamine (MEA) and 1% (v/v) GLOX was used to promote photoswitching and reduce photobleaching [39]. An inverted Nikon Ti2 Eclipse STORM 5.0 system with Perfect Focus focal plane lock was used for image acquisition. This system contains a NSTORM quadband filter, and 405, 488, 561, and 647 nm lasers and was equipped with an HP APO TIRF AC 100x/1.49 NA oil objective and ORCA-Flash4.0 SCI CMOS PLUS camera (Hamamatsu Photonics). 15,000 frames for each dye were collected at 30 ms exposure time. Localizations were identified with NIS Elements 5.0 (Nikon Instruments) and exported as tab-delimited text files.

### Clus-DoC analysis of single molecule localizations

To avoid prejudicial selection of membrane regions for analysis, we used localization data prior to cluster analysis to specify ROIs; thereby restricting our analysis to localizations that represent molecules on the nuclear envelope. EM studies demonstrated the localization of 5-LO and FLAP in the nuclear envelope after cell activation [21]. We employed Clus-DoC [31], which quantifies colocalization of individual proteins and molecules (localizations) and cluster properties. Clus-DoC allows for the user to define the number and DoC threshold.

### Unbiased cluster analysis

We implemented a modified form of the variable bandwidth mean shift (VBMS) algorithm with automatic bandwidth selection previously described [40], using the diagonal bandwidth matrix estimation method. The final bandwidth used (250 nM) was then selected from these estimates automatically, and is allowed to vary from point to point, thereby adapting to variations in scale and structure between and within datasets.

Our modifications were primarily designed to improve analysis time. Running all test bandwidths in parallel rather than in serial decreased execution time over 10-fold. Storing the STORM data in kd-trees and calculating kernel updates using points within only 4x the bandwidth of the current mean provided faster mean shift direction estimates without sacrificing accuracy. The data was preprocessed with the DBSCAN algorithm [41], with the distance cutoff set based on the 99.9^th^ percentile of nearest-neighbor distances within the dataset. Regions which were determined to be independent during this conservative DBSCAN pre-clustering stage were processed independently in parallel. Because the VBMS algorithm scales nonlinearly with the number of points, this optimization further increased processing efficiency beyond that provided by parallelization. Single isolated points were also removed at this stage, providing another ~10x increase in processing speed.

### Statistics

EIA was analyzed using one-way ANOVA followed by Bonferroni multiple comparison post-test where p < 0.05 was considered significant. Unbiased cluster property histograms were first tested for normality using Kolmogorov-Smirnov and D’Agostino-Pearson omnibus tests. After passing the normality tests (p > 0.5), the data were tested for differences between samples by one-way ANOVA followed by Bonferroni multiple comparison post-test where p < 0.05 was considered significant. A Welch’s unpaired t-test (unequal variances) was performed to determine to significance where two conditions were compared. All statistical tests were performed using Graphpad Prism 7 (Graphpad Software).

### Source code

Unbiased cluster analysis code was written for 64-bit MATLAB R2013b (MathWorks) or higher under a Windows operating system. The latest version of the source code is available via the authors’ Git repository (https://github.com/bairangie/sobermanclusters).

## Supporting Information

**S1 Fig. Strategy for analysis of the higher order assemblies of 5-LO and FLAP via unbiased cluster analysis.** RBL-2H3 mast cells were primed with anti-TNP antibody followed by crosslinking of FcεR1 by addition of TNP-BSA. Media was removed from each well and analyzed for LTC_4_. Cells were fixed, permeabilized, and prepared for molecular imaging. STORM localization lists were converted to tab-delimited text files. Image panels illustrate the process. (A) STORM image of FLAP localizations in an activated mast cell. (B) A region of interest (ROI) is drawn around the nuclear envelope FLAP localizations. (C) Localizations inside the ROI (orange) are clustered by unbiased cluster analysis. (D) The convex hull of points in a cluster (inset area from C) defines the cluster area, from which properties are calculated including number of localizations, area and density. For 5-LO, the entire nucleus and perinuclear region were included in the ROIs because 5-LO is cytoplasmic in unstimulated cells and thus the nuclear envelope is not apparent. Scale bar = 4 μm.

**S2 Fig. Frequency distributions of DoC scores for 5-LO and FLAP.** Localization data was collected by two-color dSTORM and analyzed with ClusDoC. The cells shown in Fig 2 were used to calculate DoC scores. (A) Histograms of DoC scores of all molecules for 5-LO (green) and FLAP (red) at 2min, (B) 7min, (C) 10 min.

**S3 Fig. Cluster maps for both 5-LO and FLAP.** RBL-2H3 cells were primed with anti-TNP IgE then activated with TNP-BSA for 0, 2, 5 and 10 min. Localization data was collected by two-color dSTORM and analyzed with ClusDoC. Cluster maps for 5-LO (A, green) and FLAP (B, red) from representative cells from Fig 2 over time were generated. Nonclustered localizations are colored gray.

**S4 Fig. Frequency distribution analysis of 5-LO clusters.** RBL-2H3 cells were primed with anti-TNP IgE then activated with TNP-BSA for 0, 2, 5 and 10 min and analyzed as shown S1 Fig. Cells were imaged with conventional STORM and cluster properties were analyzed with unbiased cluster analysis. (A-C) Normalized point-weighted histograms with inset bars showing mean ± SEM for (A) number of localizations, (B) cluster areas and (C) cluster densities. One-way ANOVA with Bonferroni post hoc test was performed to determine significance, indicated by ****p < 0.0005. At least 3 separate experiments collected between 10 and 30 cells.

**S5 Fig. Inhibition of cPLA_2_ and FLAP controls 5-LO and FLAP higher order assemblies.** RBL-2H3 cells were incubated with or without cPLA_2_ Inh or MK886, and then primed with anti-TNP IgE. They were then stimulated by the addition of TNP-BSA for 7 min. The cells were imaged with conventional STORM, and cluster properties were analyzed with unbiased cluster analysis. (A-F) Normalized point-weighted histograms with inset bars showing mean ± SEM for (A,D) number of localizations, (B,E) cluster areas and (C,F) cluster densities for 5-LO and FLAP, respectively. The area shaded blue represents localizations in cells primed and activated for 7 min. The solid red line represents cells incubated with cPLA_2_ Inh and primed and activated. The dotted yellow line represents cells incubated with MK886 and primed and activated. One-way ANOVA with Bonferroni post hoc test was performed to determine significance, indicated by *p < 0.05 and ***p = 0.0005. At least 3 separate experiments collected between 10 and 30 cells.

**S1 Data. Properties of clusters identified by Clus-DoC for each ROI from two-color dSTORM.**

**S2 Data. Localizations for each ROI from two-color dSTORM accepted by Clus-DoC for analysis for NT, 2 and 5 min.**

**S3 Data. Localizations for each ROI from two-color dSTORM accepted by Clus-DoC for analysis for 7 and 10 min.**

**S1 Table. Summary of clustering data for conventional STORM.**

## References

1. Galli SJ, Tsai M. IgE and mast cells in allergic disease. Nat Med. 2012;18(5):693–704. doi: 10.1038/nm.2755. PubMed PMID: 22561833; PubMed Central PMCID: PMC3597223.

2. Turner H, Kinet JP. Signalling through the high-affinity IgE receptor Fc epsilonRI. Nature. 1999;402(6760 Suppl):B24–30. Pub Med PMID: 10586892.

3. Stone KD, Prussin C, Metcalfe DD. IgE, mast cells, basophils, and eosinophils. J Allergy Clin Immunol. 2010;125(2 Suppl 2):S73–80. doi: 10.1016/j.jaci.2009.11.017. PubMed PMID: 20176269; PubMed Central PMCID: PMC2847274.

4. Scharenberg AM, Lin S, Cuenod B, Yamamura H, Kinet JP. Reconstitution of interactions between tyrosine kinases and the high affinity IgE receptor which are controlled by receptor clustering. EMBO J. 1995;14(14):3385–94. PubMed PMID: 7628439; PubMed Central PMCID: PMC394405.

5. Kinet JP. The high-affinity IgE receptor (FcεRI): from physiology to pathology. Annu Rev Immunol. 1999;17:931–72. doi: 10.1146/annurev.immunol.17.1.931. PubMed PMID: 10358778.

6. Jouvin MH, Adamczewski M, Numerof R, Letourneur O, Valle A, Kinet JP. Differential control of the tyrosine kinases Lyn and Syk by the two signaling chains of the high affinity immunoglobulin E receptor. J Biol Chem. 1994;269(8):5918–25. PubMed PMID: 8119935.

7. Phong BL, Avery L, Sumpter TL, Gorman JV, Watkins SC, Colgan JD, et al. Tim-3 enhances FcεRI-proximal signaling to modulate mast cell activation. J Exp Med. 2015;212(13):2289–304. doi: 10.1084/jem.20150388. PubMed PMID: 26598760; PubMed Central PMCID: PMC4689164.

8. Lundeen KA, Sun B, Karlsson L, Fourie AM. Leukotriene B4 receptors BLT1 and BLT2: expression and function in human and murine mast cells. J Immunol. 2006;177(5):3439–47. PubMed PMID: 16920986.

9. Weller CL, Collington SJ, Brown JK, Miller HR, Al-Kashi A, Clark P, et al. Leukotriene B4, an activation product of mast cells, is a chemoattractant for their progenitors. J Exp Med. 2005;201(12):1961–71. doi: 10.1084/jem.20042407. PubMed PMID: 15955837; PubMed Central PMCID: PMC2212026.

10. Orange RP, Valentine MD, Austen KF. Antigen-induced release of slow reacting substance of anaphylaxis (SRS-A rat) in rats prepared with homologous antibody. J Exp Med. 1968;127(4):767–82. PubMed PMID: 4384530; PubMed Central PMCID: PMC2138477.

11. Murphy RC, Hammarstrom S, Samuelsson B. Leukotriene C: a slow-reacting substance from murine mastocytoma cells. Proc Natl Acad Sci U S A. 1979;76(9):4275–9. PubMed PMID: 41240; PubMed Central PMCID: PMC411556.

12. Mencia-Huerta JM, Razin E, Ringel EW, Corey EJ, Hoover D, Austen KF, et al. Immunologic and ionophore-induced generation of leukotriene B4 from mouse bone marrow-derived mast cells. J Immunol. 1983;130(4):1885–90. PubMed PMID: 6300233.

13. Perkins JR, Diboun I, Dessailly BH, Lees JG, Orengo C. Transient protein-protein interactions: structural, functional, and network properties. Structure. 2010;18(10):1233–43. doi: 10.1016/j.str.2010.08.007. PubMed PMID: 20947012.

14. Subbotin RI, Chait BT. A pipeline for determining protein-protein interactions and proximities in the cellular milieu. Mol Cell Proteomics. 2014;13(11):2824–35. doi: 10.1074/mcp.M114.041095. PubMed PMID: 25172955; PubMed Central PMCID: PMC4223475.

15. Monachino E, Spenkelink LM, van Oijen AM. Watching cellular machinery in action, one molecule at a time. J Cell Biol. 2017;216(1):41–51. doi: 10.1083/jcb.201610025. PubMed PMID: 27979907; PubMed Central PMCID: PMC5223611.

16. Glover S, de Carvalho MS, Bayburt T, Jonas M, Chi E, Leslie CC, et al. Translocation of the 85-kDa phospholipase A2 from cytosol to the nuclear envelope in rat basophilic leukemia cells stimulated with calcium ionophore or IgE/antigen. J Biol Chem. 1995;270(25):15359–67. PubMed PMID: 7797525.

17. Gijon MA, Spencer DM, Kaiser AL, Leslie CC. Role of phosphorylation sites and the C2 domain in regulation of cytosolic phospholipase A2. J Cell Biol. 1999;145(6):1219–32. PubMed PMID: 10366595; PubMed Central PMCID: PMC2133140.

18. Brock TG, Paine R, 3rd, Peters-Golden M. Localization of 5-lipoxygenase to the nucleus of unstimulated rat basophilic leukemia cells. J Biol Chem. 1994;269(35):22059–66. PubMed PMID: 8071328.

19. Brock TG, McNish RW, Peters-Golden M. Translocation and leukotriene synthetic capacity of nuclear 5-lipoxygenase in rat basophilic leukemia cells and alveolar macrophages. J Biol Chem. 1995;270(37):21652–8. PubMed PMID: 7665580.

20. Chen XS, Naumann TA, Kurre U, Jenkins NA, Copeland NG, Funk CD. cDNA cloning, expression, mutagenesis, intracellular localization, and gene chromosomal assignment of mouse 5-lipoxygenase. J Biol Chem. 1995;270(30):17993–9. PubMed PMID: 7629107.

21. Woods JW, Coffey MJ, Brock TG, Singer, II, Peters-Golden M. 5-Lipoxygenase is located in the euchromatin of the nucleus in resting human alveolar macrophages and translocates to the nuclear envelope upon cell activation. J Clin Invest. 1995;95(5):2035–46. doi: 10.1172/JCI117889. PubMed PMID: 7738170; PubMed Central PMCID: PMC295787.

22. Mandal AK, Jones PB, Bair AM, Christmas P, Miller D, Yamin TT, et al. The nuclear membrane organization of leukotriene synthesis. Proc Natl Acad Sci U S A. 2008;105(51):20434–9. Epub 2008/12/17. doi: 10.1073/pnas.0808211106. PubMed PMID: 19075240; PubMed Central PMCID: PMC2629249.

23. Bair AM, Turman MV, Vaine CA, Panettieri RA, Jr., Soberman RJ. The nuclear membrane leukotriene synthetic complex is a signal integrator and transducer. Mol Biol Cell. 2012;23(22):4456–64. doi: 10.1091/mbc.E12-06-0489. PubMed PMID: 23015755; PubMed Central PMCID: PMC3496618.

24. Ferguson AD, McKeever BM, Xu S, Wisniewski D, Miller DK, Yamin TT, et al. Crystal structure of inhibitor-bound human 5-lipoxygenase-activating protein. Science. 2007;317(5837):510–2. doi: 10.1126/science.1144346. PubMed PMID: 17600184.

25. Christmas P, Weber BM, McKee M, Brown D, Soberman RJ. Membrane localization and topology of leukotriene C4 synthase. J Biol Chem. 2002;277(32):28902–8. Epub 2002/05/23. doi: 10.1074/jbc.M203074200. PubMed PMID: 12023288.

26. Rius M, Hummel-Eisenbeiss J, Keppler D. ATP-dependent transport of leukotrienes B4 and C4 by the multidrug resistance protein ABCC4 (MRP4). J Pharmacol Exp Ther. 2008;324(1):86–94. Epub 2007/10/26. doi: 10.1124/jpet.107.131342. PubMed PMID: 17959747.

27. Wu H. Higher-order assemblies in a new paradigm of signal transduction. Cell. 2013;153(2):287–92. Epub 2013/04/16. doi: 10.1016/j.cell.2013.03.013. PubMed PMID: 23582320; PubMed Central PMCID: PMC3687143.

28. Wu H, Fuxreiter M. The structure and dynamics of higher-order assemblies: amyloids, signalosomes, and granules. Cell. 2016;165(5):1055–66. Epub 2016/05/21. doi: 10.1016/j.cell.2016.05.004. PubMed PMID: 27203110; PubMed Central PMCID: PMC4878688.

29. Lin SC, Lo YC, Wu H. Helical assembly in the MyD88-IRAK4-IRAK2 complex in TLR/IL¬1R signalling. Nature. 2010;465(7300):885–90. Epub 2010/05/21. doi: 10.1038/nature09121. PubMed PMID: 20485341; PubMed Central PMCID: PMC2888693.

30. Wang L, Yang JK, Kabaleeswaran V, Rice AJ, Cruz AC, Park AY, et al. The Fas-FADD death domain complex structure reveals the basis of DISC assembly and disease mutations. Nat Struct Mol Biol. 2010;17(11):1324–9. doi: 10.1038/nsmb.1920. PubMed PMID: 20935634; PubMed Central PMCID: PMC2988912.

31. Pageon SV, Nicovich PR, Mollazade M, Tabarin T, Gaus K. Clus-DoC: a combined cluster detection and colocalization analysis for single-molecule localization microscopy data. Mol Biol Cell. 2016;27(22):3627–36. doi: 10.1091/mbc.E16-07-0478. PubMed PMID: 27582387; PubMed Central PMCID: PMC5221594.

32. Veatch SL, Machta BB, Shelby SA, Chiang EN, Holowka DA, Baird BA. Correlation functions quantify super-resolution images and estimate apparent clustering due to over-counting. PLoS One. 2012;7(2):e31457. doi: 10.1371/journal.pone.0031457. PubMed PMID: 22384026; PubMed Central PMCID: PMC3288038.

33. Spiess M, Hernandez-Varas P, Oddone A, Olofsson H, Blom H, Waithe D, et al. Active and inactive beta1 integrins segregate into distinct nanoclusters in focal adhesions. J Cell Biol. 2018;217(6):1929–40. Epub 2018/04/11. doi: 10.1083/jcb.201707075. PubMed PMID: 29632027; PubMed Central PMCID: PMC5987715.

34. Moore TI, Aaron J, Chew TL, Springer TA. Measuring integrin conformational change on the cell surface with super-resolution microscopy. Cell Rep. 2018;22(7):1903–12. Epub 2018/02/15. doi: 10.1016/j.celrep.2018.01.062. PubMed PMID: 29444440; PubMed Central PMCID: PMC5851489.

35. Mandal AK, Skoch J, Bacskai BJ, Hyman BT, Christmas P, Miller D, et al. The membrane organization of leukotriene synthesis. Proc Natl Acad Sci U S A. 2004;101(17):6587–92. Epub 2004/04/16. doi: 10.1073/pnas.0308523101. PubMed PMID: 15084748; PubMed Central PMCID: PMC404089.

36. Gerstmeier J, Weinigel C, Rummler S, Radmark O, Werz O, Garscha U. Time-resolved in situ assembly of the leukotriene-synthetic 5-lipoxygenase/5-lipoxygenase-activating protein complex in blood leukocytes. FASEB J. 2016;30(1):276–85. Epub 2015/09/24. doi: 10.1096/fj.15-278010. PubMed PMID: 26396238.

37. Hafner AK, Gerstmeier J, Hornig M, George S, Ball AK, Schroder M, et al. Characterization of the interaction of human 5-lipoxygenase with its activating protein FLAP. Biochim Biophys Acta. 2015;1851(11):1465–72. Epub 2015/09/04. doi: 10.1016/j.bbalip.2015.08.010. PubMed PMID: 26327594.

38. Gerstmeier J, Newcomer ME, Dennhardt S, Romp E, Fischer J, Werz O, et al. 5-Lipoxygenase-activating protein rescues activity of 5-lipoxygenase mutations that delay nuclear membrane association and disrupt product formation. FASEB J. 2016;30(5):1892–900. Epub 2016/02/05. doi: 10.1096/fj.201500210R. PubMed PMID: 26842853; PubMed Central PMCID: PMC4836370.

39. Dempsey GT. A user’s guide to localization-based super-resolution fluorescence imaging. Methods Cell Biol. 2013;114:561–92. doi: 10.1016/B978-0-12-407761-4.00024-5. PubMed PMID: 23931523.

40. Comaniciu DR, V.; Meer, P. The variable bandwidth mean shift and data-driven scale selection. IEEE Conf Pub. 2001;1:438–45. doi: 10.1109/ICCV.2001.937550.

41. Ester MK, HP.; Sander, J.; Xu, X. A density-based algorithm for discovering clusters in large spatial databases with noise. Proc Second Int Conf Knowledge Discovery Data Mining (KDD-96). 1996.

