## Supplementary figures and images for "The organization of leukotriene biosynthesis on the nuclear envelope revealed by single molecule localization microscopy and computational analyses"

### S1 Fig

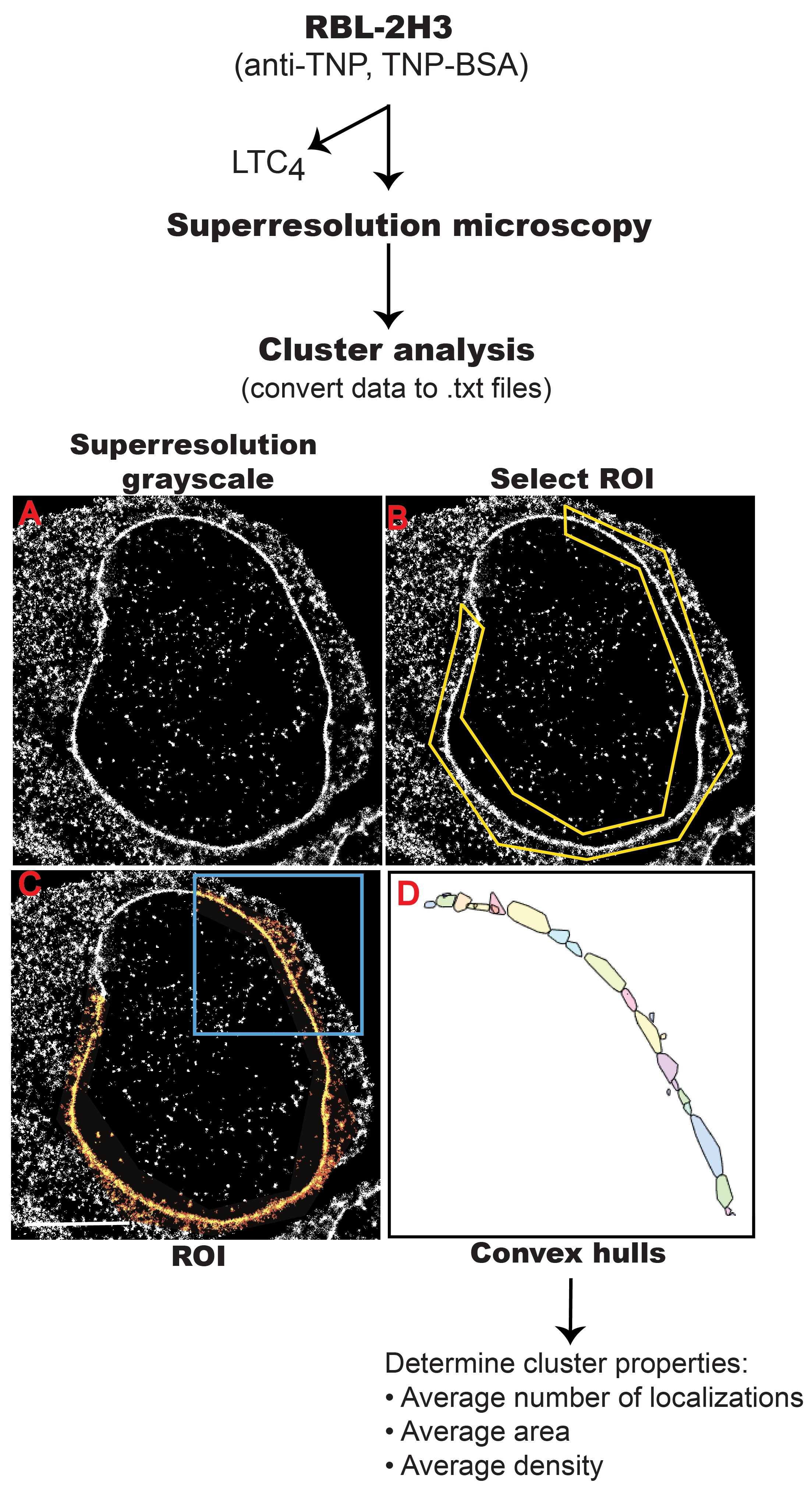

### S2 Fig

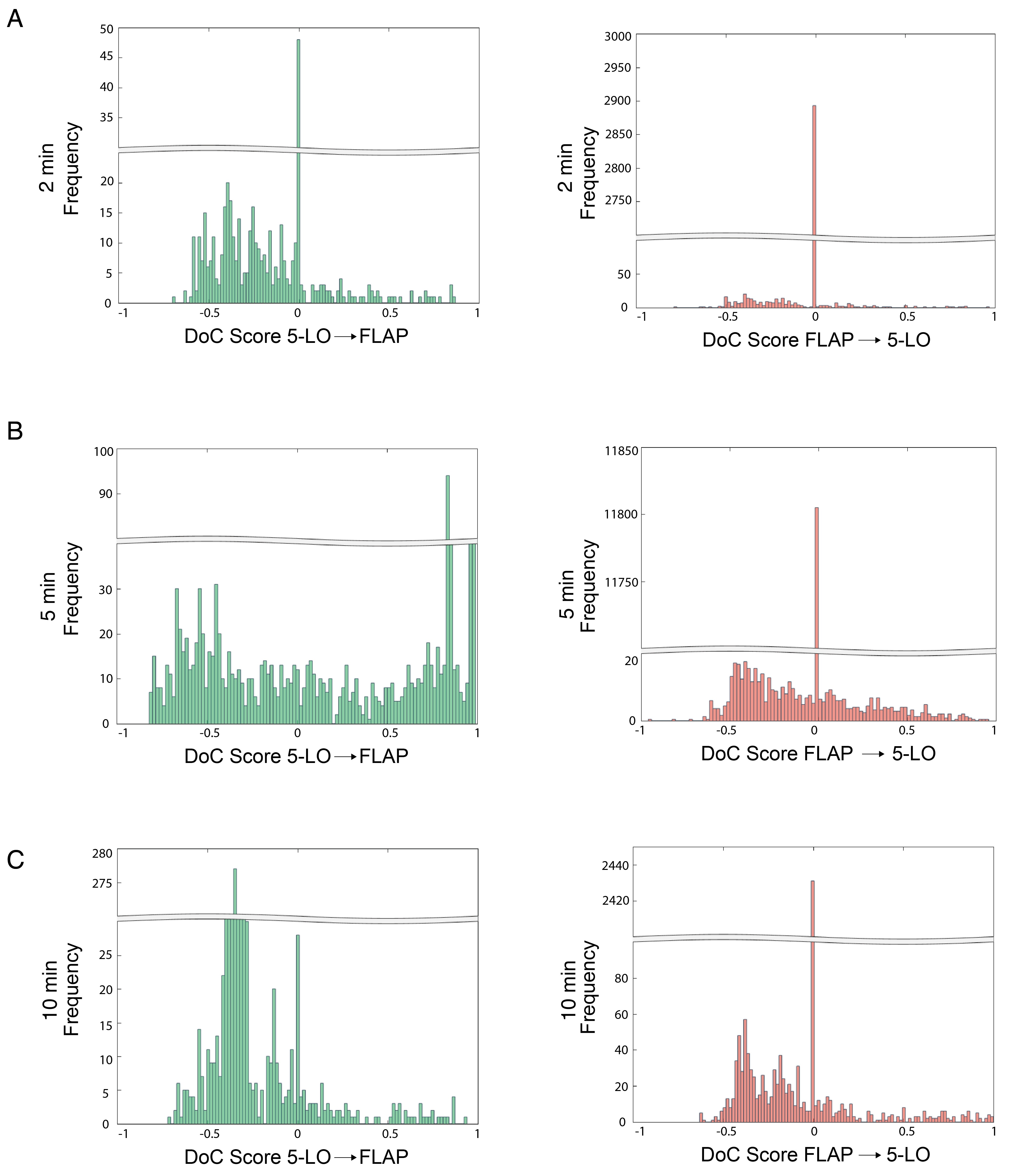

### S3 Fig

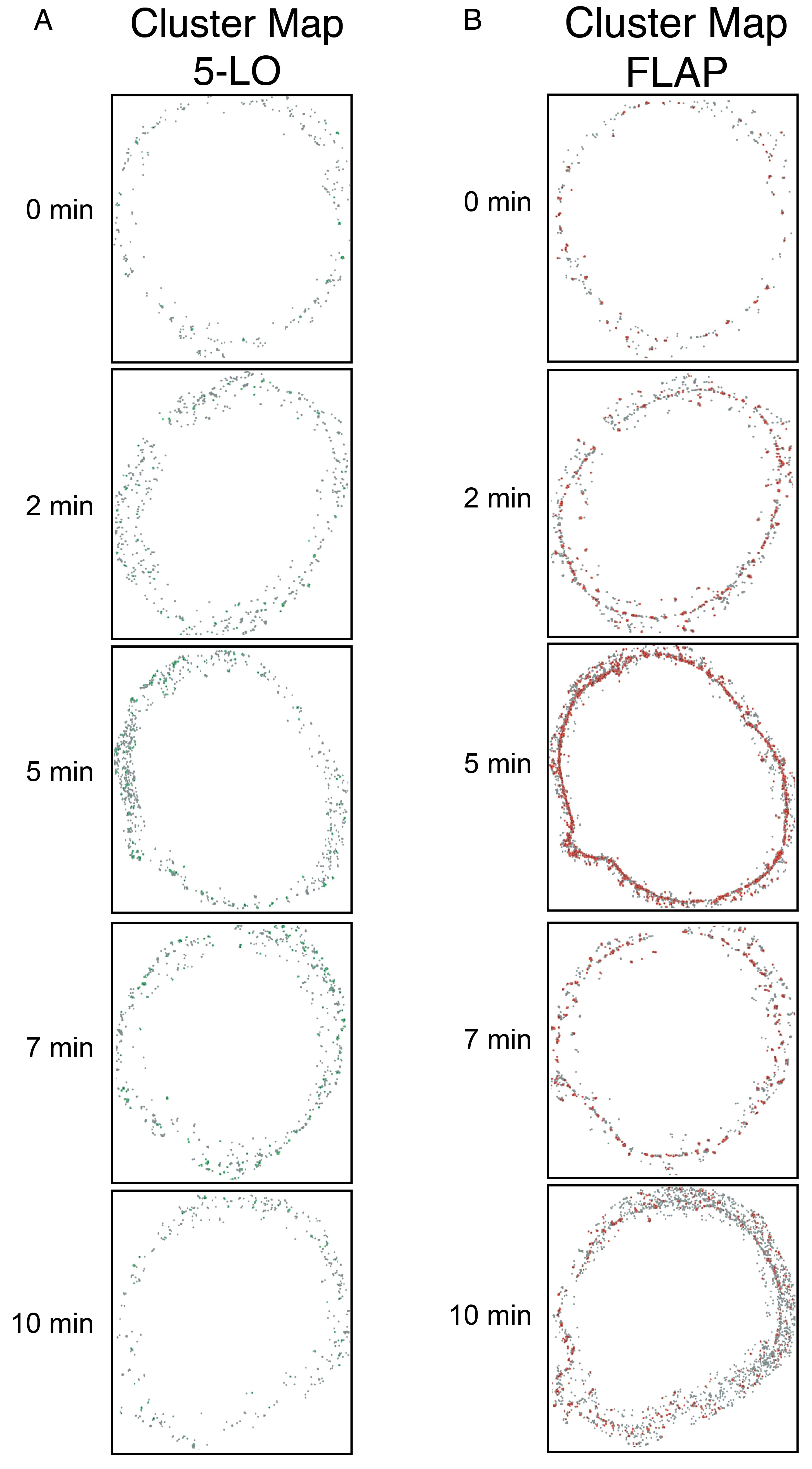

### S4 Fig

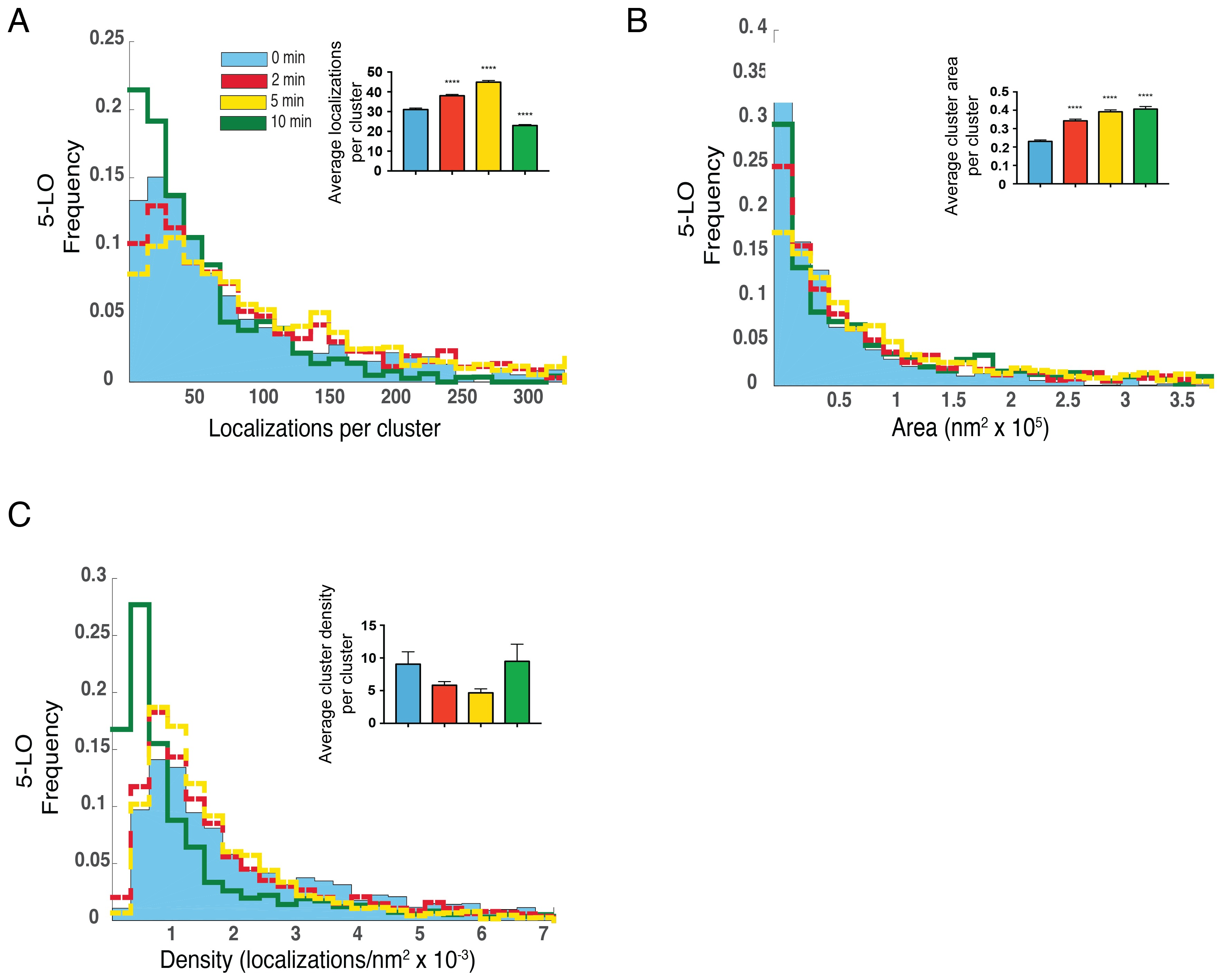

### S5 Fig

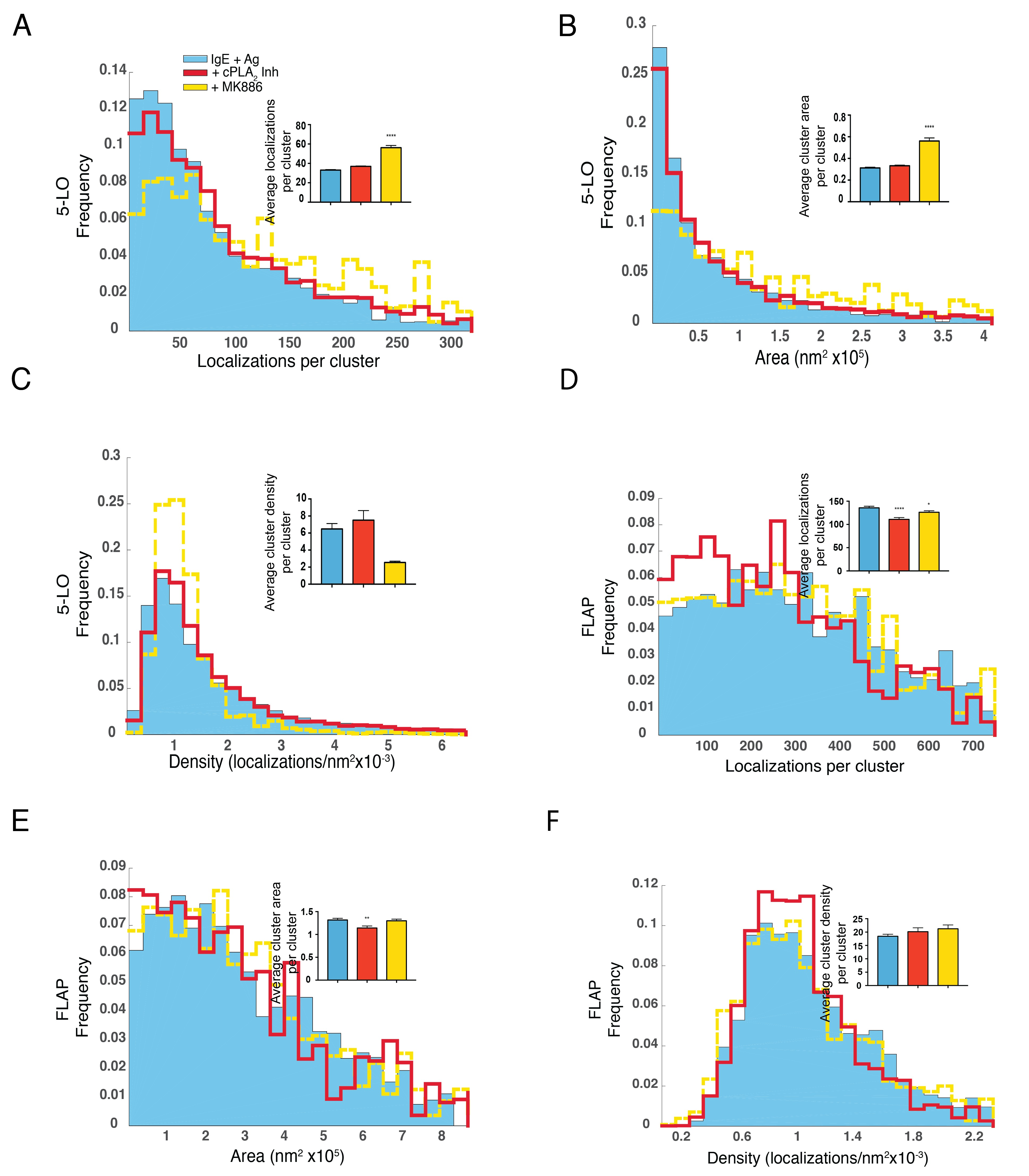
