## Supplementary material for "The organization of leukotriene biosynthesis on the nuclear envelope revealed by single molecule localization microscopy and computational analyses": S1 Table

| **Figure** | **Molecule** | **Condition** | **Localizations** | **Clusters** | **ROIs** |
| --- | --- | --- | --- | --- | --- |
| 5 | 5-LO | 0 min IgE + Ag | 113629 | 3659 | 11 |
| 5 | 5-LO | 2 min IgE + Ag | 223661 | 5885 | 16 |
| 5 | 5-LO | 5 min IgE + Ag | 247940 | 5521 | 13 |
| 5 | 5-LO | 10 min IgE + Ag | 75772 | 3298 | 7 |
| 6 | FLAP | IgE only | 128923 | 1018 | 14 |
| 6 | FLAP | 7 min IgE + Ag | 273524 | 3168 | 25 |
| S5 | 5-LO | IgE + Ag  DMSO | 283551 | 8560 | 20 |
| S5 | 5-LO | IgE + Ag + cPLA_2_ Inh | 618230 | 16713 | 42 |
| S5 | 5-LO | IgE + Ag + MK886 | 70364 | 1251 | 4 |
| S5 | FLAP | IgE + Ag  DMSO | 344535 | 2540 | 13 |
| S5 | FLAP | IgE + Ag  + cPLA_2_ Inh | 152598 | 1375 | 22 |
| S5 | FLAP | IgE + Ag  + MK886 | 337260 | 2672 | 12 |

S1 Table
